# Knockout of *tusA* facilitates flagella formation and cationic antimicrobial resistance by disrupting Fur transcriptional regulation in *Escherichia coli*

**DOI:** 10.1101/2025.06.16.659981

**Authors:** Kazuya Ishikawa, Kiho Nakata, Kazuyuki Furuta, Chikara Kaito

## Abstract

tRNA 2-thiouridine synthesizing protein A (TusA), a sulfur-carrier protein, plays a crucial role in tRNA sulfur modification. Recent studies have reported that *tusA* deficiency affects iron-sulfur (Fe-S) homeostasis and cluster formation in *Escherichia coli*; however, its association with this phenotype remains unclear. In this study, we analyzed the phenotype of *tusA*-deficient *E. coli* (Δ*tusA*) and its underlying mechanisms using RNA sequencing. We observed that *tusA* deletion disrupted the expression of genes regulated by the ferric uptake regulator Fur or Fur-regulated transcription factors (flagellar transcriptional regulators D and C [FlhDC] and fumarate and nitrate reduction regulator [Fnr]). Increased expression of *flhDC*, which is the master regulator of flagellar genes facilitated flagella formation even under conditions in which the wild-type formed few flagella. Additionally, Δ*tusA* was resistant to cationic antibacterial agents, such as cetyltrimethylammonium bromide, cetylpyridinium chloride, and protamine sulfate. This resistance is associated with the increased expression of *ompX* or *ompF*, regulated by Fur and Fnr, respectively. Notably, both enhanced flagella formation and resistance to cationic antibacterial agents caused by *tusA* deletion were abolished in the *fur*-deficient background. These findings indicate that impaired expression of the *fur* regulon, possibly because of impaired Fe-S cluster formation, induces multiple phenotypic alterations in Δ*tusA*.

**IMPORTANCE:** TusA is a sulfur carrier protein involved in tRNA sulfur modification, and its effect on translation has been studied. Recent studies have reported that *tusA* deficiency affects Fe-S homeostasis and cluster formation in *Escherichia coli*; however, its association with the phenotype remains unclear. Based on RNA sequencing, we indicated that the altered gene expression in Δ*tusA* resulted from the disruption of fur regulation that is controlled by Fe-S clusters. We further demonstrated that enhanced flagella formation and resistance to cationic drugs were mediated by Fur-dependent gene expression alterations. Our data indicate that the regulation of sulfur allocation for tRNA modification by TusA affects the global gene expression in bacteria.

## INTRODUCTION

tRNA 2-thiouridine synthesizing protein A (TusA) is a sulfur-carrier protein that functions as a sulfur transferase in molybdenum coenzyme biosynthesis and tRNA modification (1, 2). TusA accepts sulfur from cysteine through the function of cysteine desulfurase IscS and transfers it to the TusBCD complex, TusE, and MnmA. This results in the thiolation of uridine at the wobble position 34 of the tRNAs of Lys, Gln, and Glu (mnm5s2U modification) (3, 4). This thiolation rearranges the tRNA structure to achieve high accuracy and efficiency in protein translation (5).

*TusA*-deficient (Δ*tusA*) *E. coli* exhibits the following phenotypes—reduced normal cell growth (6, 7), phage resistance because of altered frameshift efficiency (7), and reduced translation efficiency of the transcriptional regulator factor for inversion stimulation (Fis) and the sigma factor RNA polymerase sigma S factor (RpoS) (8). Additionally, microarray analysis demonstrated that in Δ*tusA*, the expression of genes associated with molybdenum cofactor biosynthesis and molybdoenzymes was increased, whereas the expression of genes involved in iron-sulfur (Fe-S) cluster formation was decreased (9). Fe-S clusters are essential cofactors primarily found in proteins that mediate electron transfer, redox reactions, and substrate binding/activation (10). tRNA thiolation and iron-sulfur cluster biosynthesis are interdependent because both require cysteine as a sulfur source and rely on IscS as a sulfur transferase (11). Recently, *tusA* deletion was shown to reduce the translation of ferric uptake regulator (Fur), resulting in reduced levels of Fe-S cluster formation in *E. coli* (9). Therefore, Δ*tusA E. coli* exhibits pleiotropic phenotypes because *tusA* is involved in translating multiple transcriptional regulators and forming Fe-S clusters. However, the association between the Δ*tusA* phenotype and underlying mechanisms remains poorly understood.

Fur is a transcription factor that regulates the expression of genes primarily associated with iron uptake and Fe-S cluster biogenesis. It is activated by binding to Fe^2+^ and inhibited by binding to Fe-S cluster proteins, thereby maintaining the intracellular balance of Fe-S clusters and iron-sulfur homeostasis (12, 13). Additionally, Fur regulates the expression of fumarate and nitrate reduction regulator (Fnr), an Fe-cluster-containing oxygen-responsive transcription factor (14), and flagellar transcriptional regulators D and C (FlhDC), a transcription factor (15). Therefore, Fur plays a crucial role in bacterial environmental adaptation, including the regulation of genes other than those associated with iron and Fe-S clusters (16).

In this study, we assessed the phenotypes of Δ*tusA E. coli* and its underlying molecular mechanisms using RNA sequencing. We observed that over half of the significantly upregulated genes in Δ*tusA* were regulated by Fur, FlhDC, or Fnr—the latter two of which are regulated by Fur. These transcription factors govern the genes involved in flagella formation and cationic drug resistance, indicating that some pleiotropic phenotypes of Δ*tusA* can be explained by the abnormal regulation of Fur expression.

## RESULTS

### RNA expression analysis of *ΔtusA E. coli*

To assess the role of *tusA* in *E. coli*, we performed RNA sequencing of Δ*tusA* in the exponential phase under aerobic conditions. Compared to the wild type, 131 genes were upregulated over 4-fold, and 200 genes were downregulated to less than one-quarter (Fig. 1A). Gene ontology analysis revealed significant enrichment of flagella formation and chemotaxis genes upregulated in Δ*tusA* (Fig. 1B), with most of the flagella formation-related genes increased over 32-fold (Fig. 1A, red dots). In contrast, genes downregulated in Δ*tusA* were enriched in metabolic degradation pathways (Fig. 1C). Numerous genes with significantly altered expression levels in Δ*tusA* were regulated by Fur, FlhDC, Fnr, leucine-responsive regulatory protein (Lrp), and Fis, whose expressions are repressed by Fur; therefore, we quantified these genes. Over half of the upregulated genes in Δ*tusA* were regulated by Fur, FlhDC, and Fnr (Fig. 1D), indicating that the regulation of expression by Fur was disrupted. The expression levels of Fur, FlhD, FlhC, and Fnr in Δ*tusA* were 1.12-, 5.05-, 4.29-, and 1.67-fold higher, respectively. This indicates that Fur-mediated suppression of FlhDC and Fnr expression is weakened in Δ*tusA*, independent of the *fur* expression level. In contrast, genes with reduced expression in Δ*tusA* were regulated by Fur, Fnr, Lrp, and Fis at similar rates. Notably, among the Fur-regulated genes, approximately half were repressed and half were activated by Fur in the wild-type. The presence of genes whose regulation by Fur is reversed indicates that the deletion of *tusA* does not simply inactivate Fur but disrupts transcriptional regulation by Fur.

**Fig 1.**
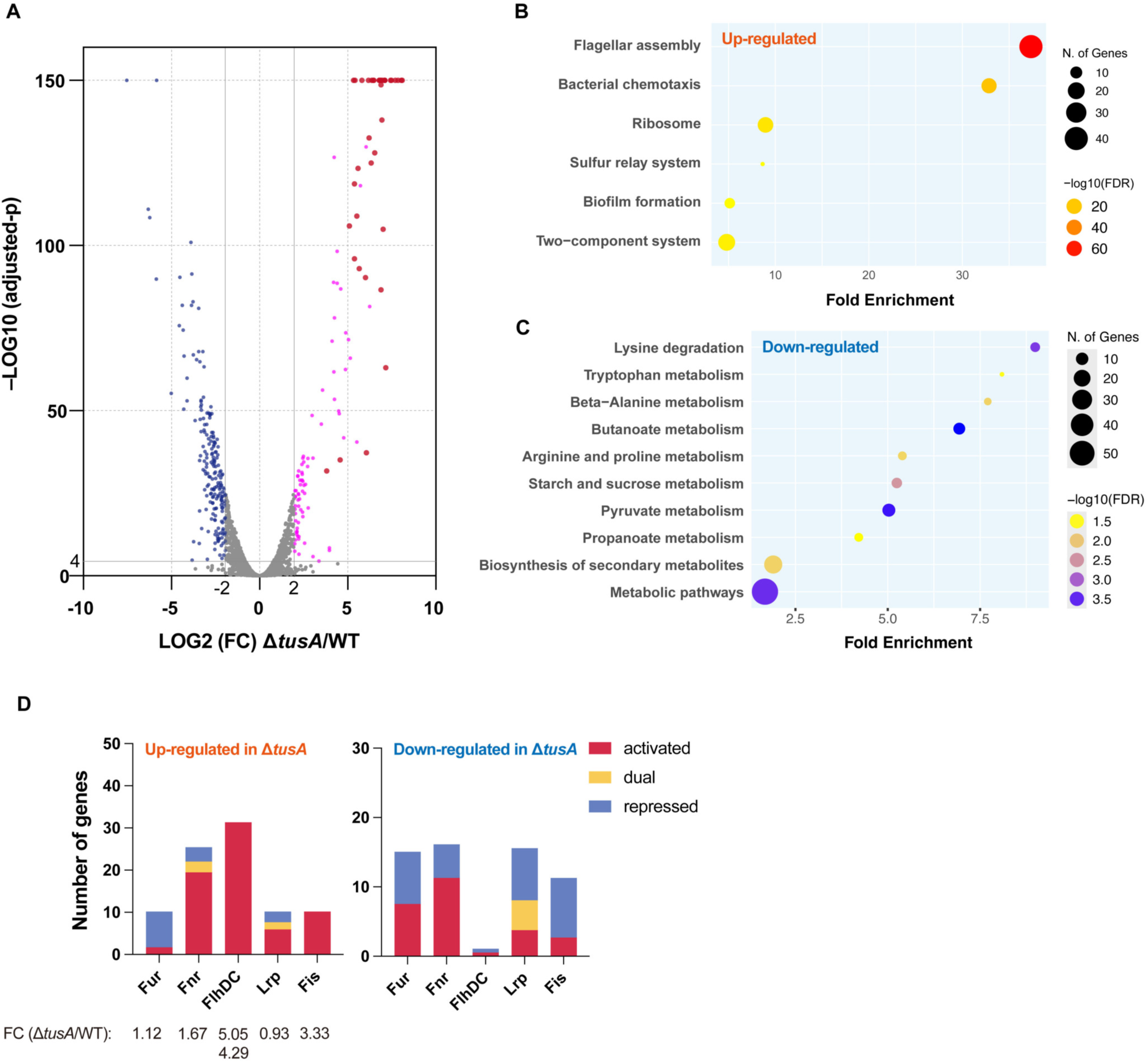
Expression of genes downstream of the ferric uptake regulator (Fur) is increased in the tRNA 2-thiouridine synthesizing protein A-deficient mutant (Δ*tusA*). (**A**) Volcano plot of RNA sequencing data. The x-axis represents the Log_2_ fold change (FC) of each gene in Δ*tusA* relative to BW25113 (wild-type [WT]). The y-axis represents the –Log_10_ adjusted p-value of each gene. Genes that demonstrated −Log_10_(adjusted p-value) ≥ 4 and Log_2_(FC) ≥ 2 or ≤ −2 are displayed in magenta and blue, respectively. The red plots indicate flagella-related genes whose expression is increased by over 32-fold. (**B, C**) Gene ontology enrichment analysis. (**B**) Enrichment of upregulated genes [-Log_10_(adjusted p-value) ≥ 4, Log_2_(FC) ≥ 2]. (**C**) Enrichment of downregulated genes [-Log_10_(adjusted p-value) ≥ 4, Log_2_(FC) ≤ 2]. **(D)** The number of genes regulated by transcription factors (Fur), fumarate and nitrate reduction regulator (Fnr), flagellar transcriptional regulators D and C (FlhDC), leucine-responsive regulatory protein (Lrp), and factor for inversion stimulation (Fis) in up- or down-regulated genes. The numbers under each transcription factor indicate the expression fold change (Δ*tusA*/WT). FlhDC demonstrates the expression fold change value of FlhC below FlhD.

### Δ*tusA* exhibits increased flagella formation

RNA sequencing revealed a significant upregulation of flagella formation-related genes in Δ*tusA*; therefore, we examined flagella formation using immunofluorescence. Consistent with the salt-induced suppression of flagella-related gene expression in BW25113 (17), flagella formation was suppressed in the wild-type BW25113 strain cultured in Luria-Bertani (LB) medium. However, excessive flagella formation was observed in Δ*tusA* in LB medium (Fig. 2A and B), and this increase was suppressed in the *fur*-deficient background. Even under low-salt conditions, where flagella formation is frequently observed in BW25113 (17), Δ*tusA* formed flagella at a higher frequency than that of the wild-type (Fig. 2A and B). Increased flagella formation was suppressed in the *fur*-deficient background, similar to that observed in LB medium. In either medium, Δ*tusA* did not form flagella in a *flhDC*-deficient background, which is a master regulator of flagellar genes. These results indicate that Fur and FlhDC regulate the increased flagella formation in Δ*tusA* at the transcriptional level. Although *flhDC* expression is repressed by Fur, neither the *fur*-deficient mutant (Δ*fur*) nor the double-deficient mutant (Δ*tusA*Δ*fur*) demonstrated increased flagella formation like Δ*tusA*. This indicates that FlhDC was abnormally activated by Fur in Δ*tusA*.

**Fig 2.**
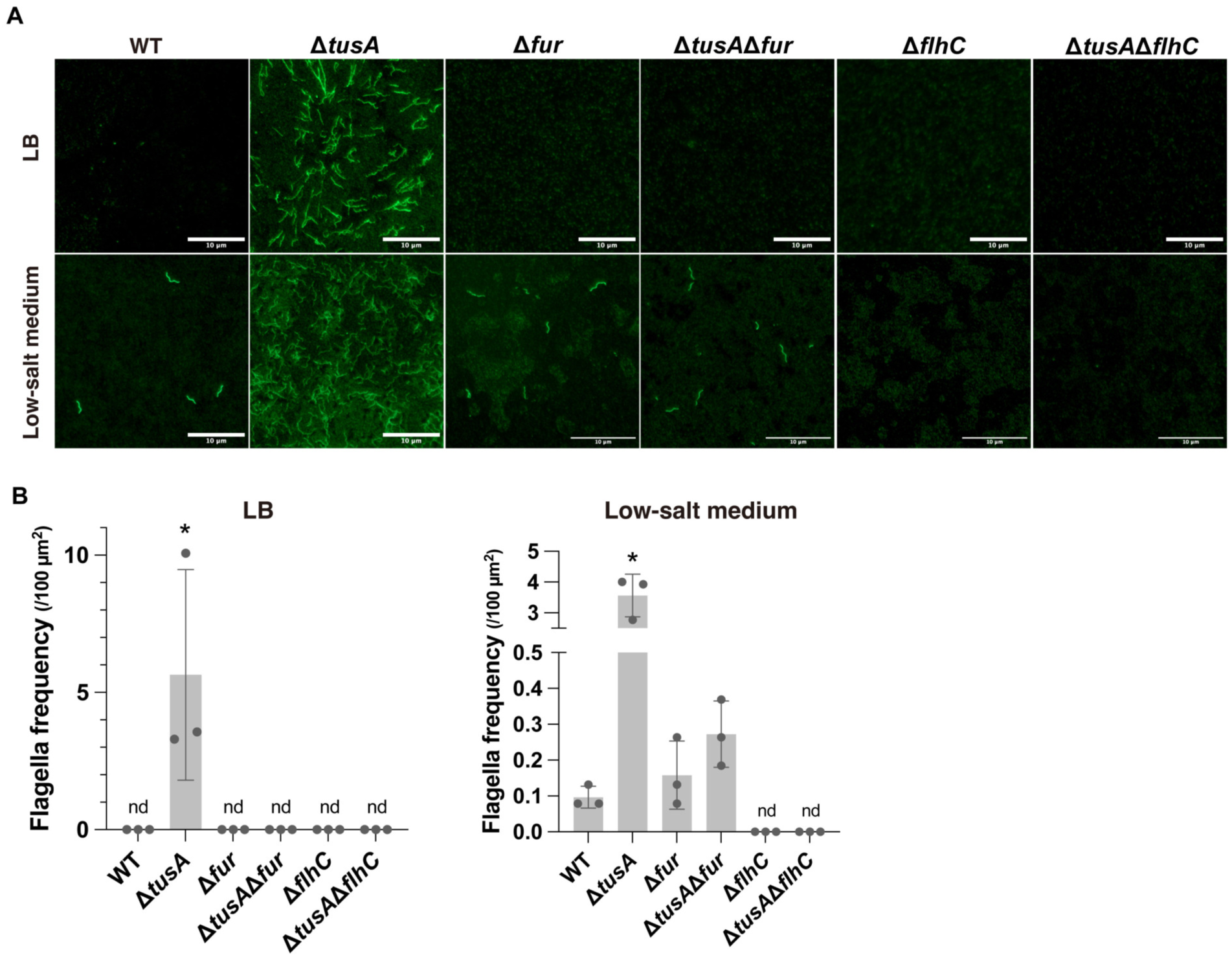
Deletion of *tusA* facilitates flagella formation in an Fur-dependent manner. (**A**) Typical images of immunofluorescence using an anti-flagellin antibody. Bacteria cultures (upper panels, LB; lower panels, low-salt medium) of BW25113 (WT) and deletion mutants (Δ*tusA*, Δ*fur*, Δ*tusA*Δ*fur*, Δ*flhC*, and Δ*tusA*Δ*flhC*) are visualized by fluorescence microscopy. (**B**) Flagella formation frequency. Flagellar frequency per area is measured using randomly acquired images. Data are presented as the means ± standard deviation; asterisks indicate significant differences, *n* = 3, *P* < 0.05, using Tukey’s multiple comparisons test.

### Deletion of *tusA* disrupts swimming motility

We assessed swimming motility because flagella formation was enhanced in Δ*tusA*. In the LB medium, slight swimming motility was observed only in Δ*tusA* (Fig. 3A). This was inconsistent with the formation of excessive flagella (Fig. 2A), indicating that Δ*tusA* had a defect in flagellar motility. Under low-salt conditions, the wild-type exhibited migration, and Δ*tusA* exhibited slightly stronger migration than that of the wild-type (Fig. 3B). Δ*fur* exhibited significantly stronger swimming motility than that of the wild-type, but the double-deficient mutant (Δ*tusA*Δ*fur*) was almost unable to migrate. The frequency of flagella formation was similar between Δ*fur* and Δ*tusA*Δ*fur* (Fig. 2), indicating that *tusA* deficiency disrupts flagellar motility through an Fur-independent pathway. In either medium, the double-deficient mutant (Δ*tusA*Δ*flhC*) did not exhibit swimming motility. These data indicate that *tusA* deficiency increased the frequency of flagella formation in an Fur-dependent manner but reduced flagellar motility in an Fur-independent manner.

**Fig 3.**
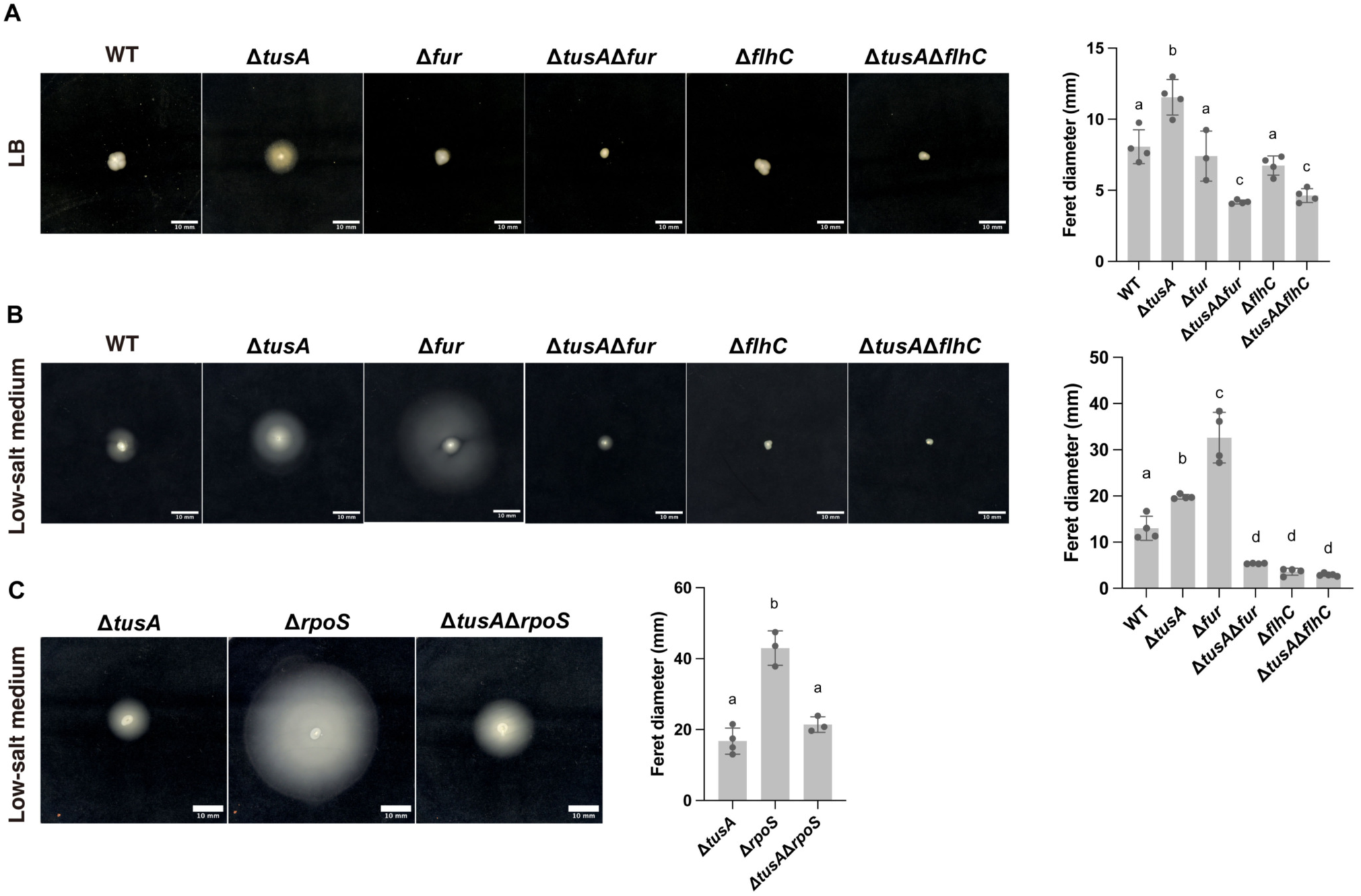
Deletion of *fur* affects the swimming motility of Δ*tusA*. (**A, B**) Images demonstrating swimming motility of BW25113 (WT) and deletion mutants (Δ*tusA*, Δ*fur*, Δ*tusA*Δ*fur*, Δ*flhC*, and Δ*tusA*Δ*flhC*) in LB (**A**) or low-salt medium (**B**) with 0.25 % agar. The graphs demonstrate the quantitative results of swimming halo. Data are presented as the means ± standard deviation (SD); different letters indicate significant differences within the same medium, *n* = 3 or 4, *P* < 0.05, using Tukey’s multiple comparisons test. (**C**) Images demonstrate swimming motility of deletion mutants (Δ*tusA*, Δ*rpoS*, and Δ*tusA*Δ*rpoS*) in low-salt medium with 0.25 % agar. Data are presented as the means ± SD; different letters indicate significant differences, *n* = 3 or 4, *P* < 0.05, using Tukey’s multiple comparisons test.

*TusA* deletion affects *rpoS* translation efficiency (8), and *rpoS*-deficient mutant (Δ*rpoS*) exhibits increased swimming motility (18). Therefore, we hypothesized that *rpoS* may alter the swimming motility of Δ*tusA* and compared the motility of the double-deficient mutant (Δ*tusA*Δ*rpoS*) with that of each single-deficient strain. Δ*rpoS* exhibited enhanced swimming motility, consistent with previous studies (Fig. 2C). Swimming motility in Δ*tusA*Δ*rpoS* was reduced, reaching almost the same level as that in Δ*tusA*. These results indicate that *tusA* deficiency reduces motility through an RpoS-independent pathway.

### Deletion of *tusA* confers resistance to cationic antimicrobials

Assessing the antimicrobial susceptibility of Δ*tusA* revealed increased resistance to cationic drugs, including cetyltrimethylammonium bromide (CTAB) and cetylpyridinium chloride (CPC), and the polycationic peptide protamine sulfate (PS) (Fig. 4A). We assessed the susceptibility of the wild-type strain to the anionic and non-ionic detergents, sodium dodecyl sulfate (SDS) and Triton X-100, respectively. However, the wild-type did not demonstrate a reduced colony formation even at a high concentration of 4 %, and therefore it was not possible to determine whether Δ*tusA* was resistant to these detergents (Fig. 4B). Because excessive flagella in Δ*tusA* may protect cells from cationic drugs, we assessed the effects of CTAB, CPC, and PS on Δ*tusA*Δ*flhC* (Fig. 4C). The drug sensitivity of Δ*tusA*Δ*flhC* to CTAB, CPC, and PS was not significantly different from that of Δ*tusA* (Fig. 4C), indicating that resistance to cationic antimicrobials was independent of flagella formation. Therefore, we assessed Δ*tusA*Δ*fur* sensitivity to cationic drugs. The resistance observed in Δ*tusA* was suppressed in the Δ*fur* background (Fig. 4D), indicating that the resistance of Δ*tusA* to cationic antimicrobials is caused by *fur*-dependent factors other than flagella formation.

**Fig 4.**
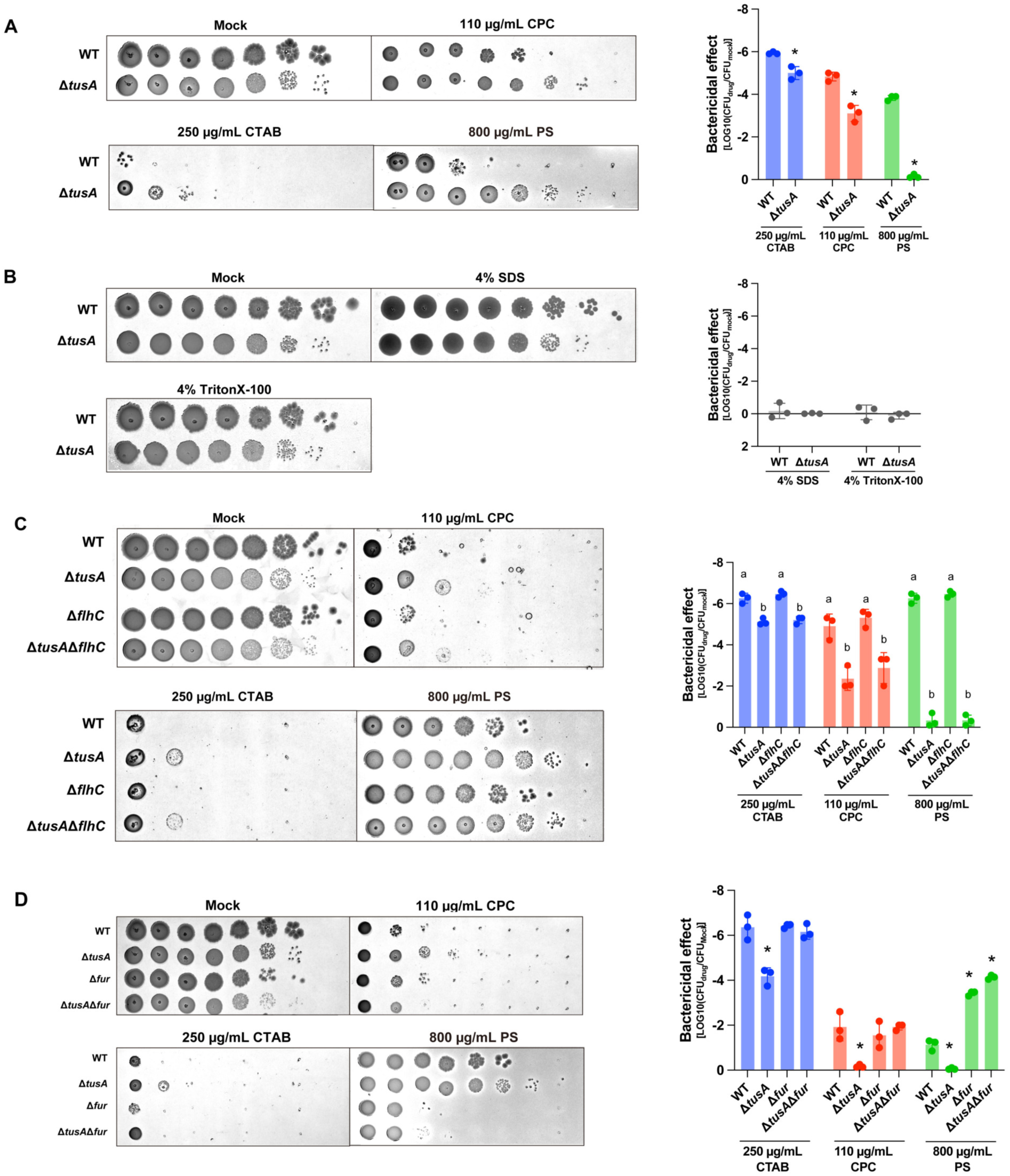
Δ*tusA* is resistant to cationic antimicrobials. Bacterial cultures are diluted 10-fold and spotted onto LB agar or LB agar with cationic antimicrobials. (**A**) Bacterial cultures dilutions of BW25113 (wild-type [WT]) and Δ*tusA* are spotted onto LB agar or LB agar with cetyltrimethylammonium bromide (CTAB), cetylpyridinium chloride (CPC), and protamine sulfate (PS). (**B**) Bacterial culture dilutions of WT and Δ*tusA* are spotted onto LB agar or LB agar with sodium dodecyl sulfate or Triton X-100. (**C**) Bacterial cultures of WT and the deletion mutants (Δ*tusA*, Δ*flhC*, and Δ*tusA*Δ*flhC*) are spotted onto LB agar or LB agar with CTAB, CPC, and PS. The antibacterial activity is demonstrated as the Log10 value of colony forming unit (CFU) (drug)/CFU (mock). Data are presented as the means ± standard deviation (SD); different letters indicate significant differences, *n* = 3, *P* < 0.05, using Tukey’s multiple comparisons test. (**D**) Bacterial cultures of WT and the deletion mutants (Δ*tusA*, Δ*fur*, and Δ*tusA*Δ*fur*) are spotted onto LB agar or LB agar with CTAB, CPC, and PS. The antibacterial activities of Δ*tusA*, Δ*fur*, and Δ*tusA*Δ*fur* are demonstrated as the Log_10_ value of CFU (drug)/CFU (mock). Data are presented as the means ± SD; different letters indicate significant differences within the same medium, *n* = 3, *P* < 0.05, using Tukey’s multiple comparisons test.

Among the genes with altered expression in Δ*tusA* that were regulated by Fur, the outer membrane β-barrel porins outer membrane protein F (*ompF*), outer membrane protein X (*ompX*), and outer membrane protein W (*ompW*) were predicted to be involved in CTAB resistance (19, 20). In Δ*tusA*, the expression levels of *ompF* and *ompX* were 2.3- and 1.8-fold higher, respectively, whereas the expression level of *ompW* was 3.0-fold lower than that in the wild-type (Fig. 5A). *OmpF* is an Fur-regulated outer membrane porin that forms a cation-selective pore to permeate molecules < 600 kDa (21). O*mpX* expression is regulated by Fnr, which is in turn regulated by Fur and is involved in cell adhesion and SDS resistance (22). *OmpW* is regulated by both Fur and Fnr and upregulated in response to antibiotic treatment (23). Therefore, we assessed whether Δ*tusA* resistance to cationic antimicrobials was observed in *ompF*-, *ompX*-, and *ompW*-deficient backgrounds. The resistance of Δ*tusA* to CTAB and CPC was suppressed in the Δ*ompF* and Δ*ompX* backgrounds, respectively (Fig. 5B and C). Additionally, the resistance of Δ*tusA* to PS was suppressed in the Δ*ompX* background. However, the resistance of Δ*tusA* to the three cationic antimicrobials was not significantly altered in the Δ*ompW* background. These results indicated that the resistance of Δ*tusA* to cationic antimicrobials was because of the enhanced expression of *ompF* and *ompX*.

**Fig 5.**
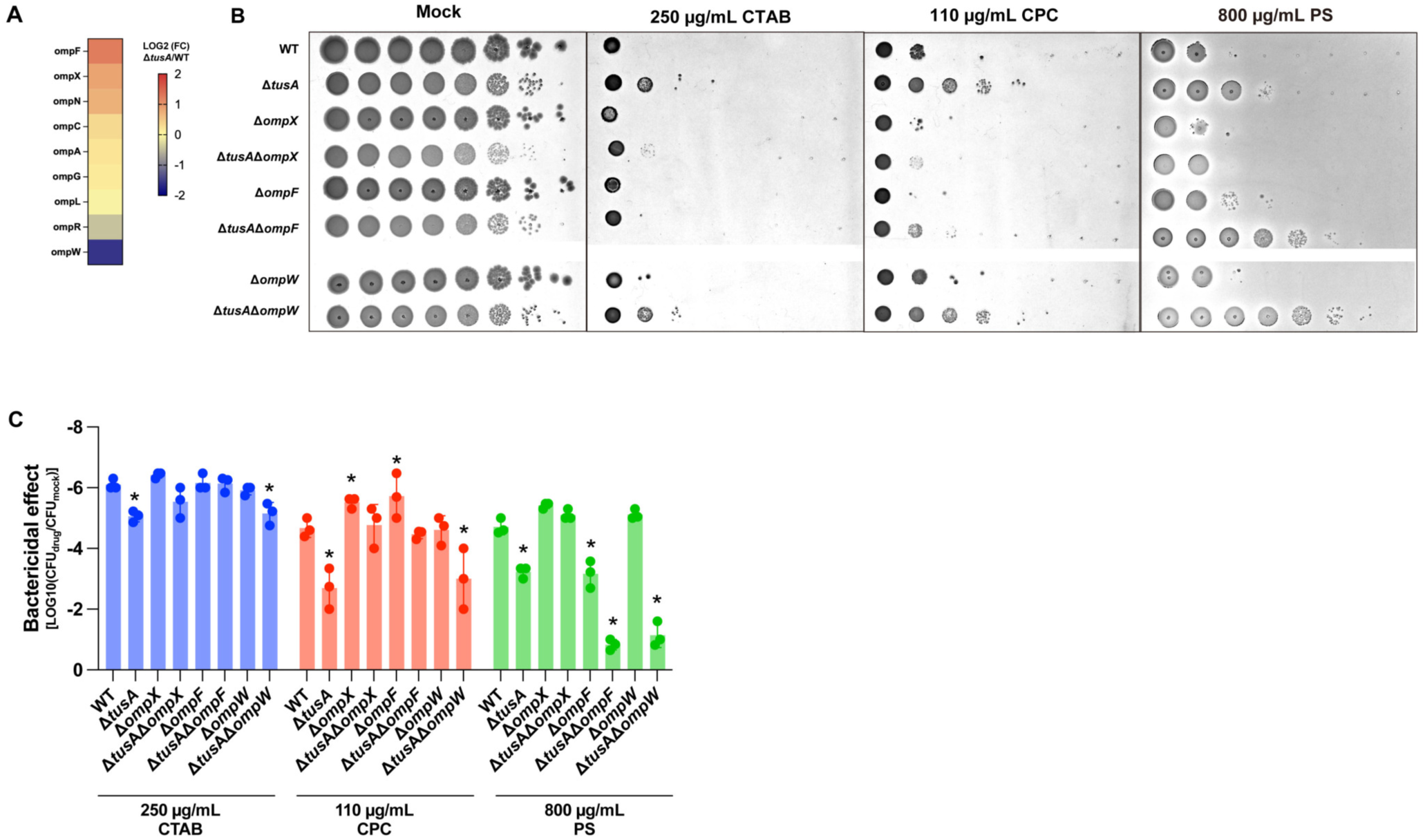
Cationic antimicrobial resistance in Δ*tusA* is dependent on *ompF* and *ompX*, which are controlled by Fur. (**A**) Expression alterations in omp genes in Δ*tusA*. The same RNA sequence data as in Fig. 1 is used. (**B**) Bacterial cultures of wild-type and the deletion mutants (Δ*tusA*, Δ*ompX*, Δ*tusA*Δ*ompX* Δ*ompF*, Δ*tusA*Δ*ompF,* Δ*ompW*, and Δ*tusA*Δ*ompW*) are spotted onto LB agar or LB agar with cetyltrimethylammonium bromide, cetylpyridinium chloride, and protamine sulfate. (**C**) The antibacterial activities of Δ*tusA*, Δ*ompX*, Δ*tusA*Δ*ompX* Δ*ompF*, Δ*tusA*Δ*ompF,* Δ*ompW*, and Δ*tusA*Δ*ompW* are demonstrated as the Log_10_ value of colony forming unit (CFU) (drug)/CFU (mock). Data are presented as the means ± standard deviation; different letters indicate significant differences within the same medium, *n* = 3, *P* < 0.05, using Tukey’s multiple comparisons test.

## DISCUSSION

RNA sequencing revealed that numerous genes whose expression was altered by *tusA* deletion belonged to the Fur regulon. Additionally, we demonstrated that two previously unidentified Δ*tusA* phenotypes, increased flagella formation with reduced motility and cationic drug resistance, depended on Fur regulation (Fig. 6). TusA, a tRNA mnm^5^s^2^U modification enzyme, has been extensively studied for its effects on translation (8, 9). However, the pleiotropic phenotype of Δ*tusA* could not be fully explained by translation deficiency alone. Our results indicate that altered expression of the Fur regulon contributes significantly to the pleiotropic phenotype of Δ*tusA*.

**Fig 6.**
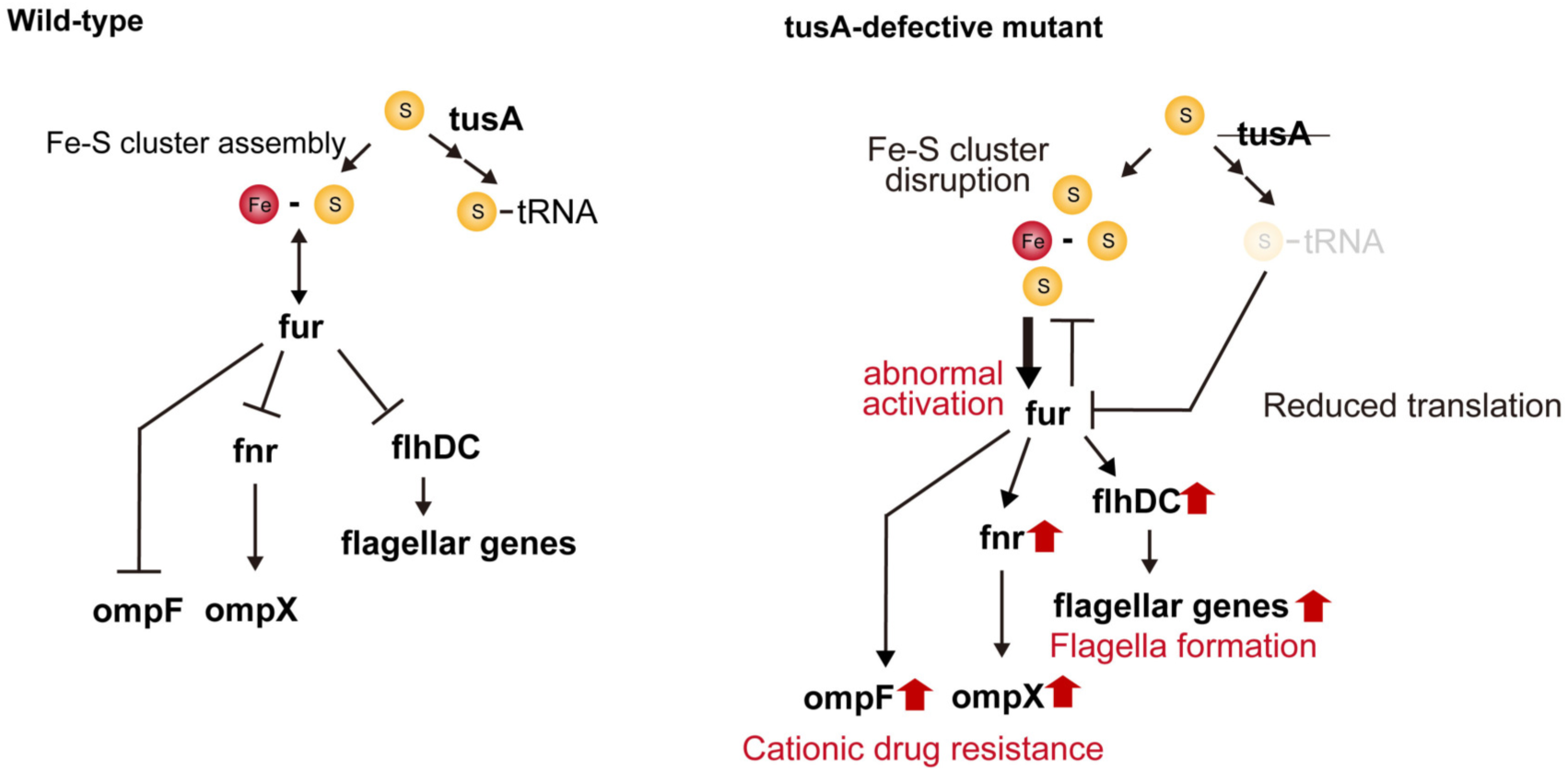
A model in which Δ*tusA* overproduces flagella with reduced motility and becomes resistant to cationic drugs.

Numerous genes with altered expression in Δ*tusA* were directly or indirectly regulated by Fur, including the *fnr* and *flhDC* regulons. Genes upregulated and downregulated by Fur in the wild-type were equally included among the genes downregulated in Δ*tusA* (Fig. 1D). This indicates that, rather than a simple inactivation of Fur, an abnormality in its transcriptional regulation occurred, such as the reversal or enhancement of gene expression patterns. This idea is supported by the observation that Δ*tusA* exhibited enhanced flagella formation (Fig. 2) and cationic drug resistance (Fig. 4 and Fig. 5) in an *fur*-dependent manner, although Δ*fur* did not exhibit these phenotypes. If Fur inactivation was responsible for the Δ*tusA* phenotype, similar phenotypes should be observed in Δ*fur*.

Although *fur* mRNA levels were slightly altered by *tusA* deletion (Fig. 1D), the translation efficiency of *fur* mRNA was reduced in Δ*tusA* (8). However, a reduction in Fur protein levels cannot fully explain the altered gene expression patterns, flagella formation, and cationic drug resistance in Δ*tusA*. Therefore, we believe that in Δ*tusA*, the abnormal transcriptional regulation by Fur results from the inhibition of Fe-S cluster formation owing to the disrupted Fe-S homeostasis. Because both tRNA thiolation and Fe-S cluster biosynthesis require sulfur atoms transferred by IscS from cysteine, they are tightly coordinated (11). Therefore, *tusA* deletion is predicted to disrupt sulfur allocation and affect Fe-S cluster biosynthesis.

The primary types of Fe-S clusters are cubic [4Fe-4S] and rhombic [2Fe-2S] (24). T*usA* deletion decreases the Fe-S cluster [4Fe-4S] levels and increases the intracellular free Fe^2+^ concentration (9). Fur is inactivated by binding to [2Fe-2S] and activated by binding to Fe^2+^ (12, 13). We demonstrated that the expression levels of the *sufABCDSE* and *iscRSUA* operons were reduced 9- and 2-fold, respectively, in Δ*tusA* compared to those in the wild-type (Table. S1). *E. coli* lacking the Isc/Suf pathway loses [4Fe-4S] but can synthesize [2Fe-2S] (12, 25). These findings indicate that *tusA* deletion disrupts sulfur allocation and affects the function or expression of *Isc*/*Suf*, thereby significantly reducing Fe-S cluster levels other than [2Fe-2S] and increasing the intracellular free Fe^2+^ concentration. This may result in the simultaneous occurrence of Fur suppression and activation owing to an increase in the relative amount of [2Fe-2S] and free Fe^2+^, respectively, thereby disrupting Fur regulation.

Swimming motility varies significantly based on the *E. coli* strain, and BW25113 has low swimming motility, specifically in the presence of salt, owing to the suppression of *flh* operon expression by the insertion of CP4-57 prophage genes (17). However, Δ*tusA* exhibited a significant increase in flagella formation in LB medium (Fig. 2). This indicates that *tusA* deficiency facilitates the expression of *flh* operon through Fur, overcoming the repression of prophage genes. Notably, flagella formed in Δ*tusA* appeared to be weakly motile (Fig. 2 and Fig. 3). In the *fur*-deficient background, *tusA* deletion had little effect on flagella formation but significantly affected swimming motility (Fig. 2 and Fig. 3). These data indicate that an Fur-independent pathway affects flagellar motility in Δ*tusA*. Flagellar proteins and factors associated with chemotaxis were collectively upregulated in Δ*tusA* (Fig. 1B). Additionally, the expression of *tsr*, *trg*, *tar*, *tap*, and *aer*, which are chemotaxis sensors regulated by Fnr or Fis but not by FlhDC, was upregulated (Table S1). Although *fnr* is regulated by Fur, Fnr itself has an Fe-S cluster. Disruption of Fe-S cluster formation may affect Fnr activity in an Fur-independent manner. Simultaneous overexpression of different chemotaxis receptors may result in excessive signaling input, affecting swimming motility.

*Omp* genes encode outer membrane porins involved in the permeation of various substances and contribute to drug resistance (26). OmpF is a highly expressed protein that generally forms pores for molecules of ≤ 600 Da (21, 27). OmpF pores serve as pathways for various drugs, and *ompF* deficiency suppresses the cellular uptake of antibiotics, such as β-lactams and quinolones, conferring antibiotic resistance (28–30). Although the role of OmpX in substance permeation is largely unknown, *E. coli* lacking *ompX* reduced virulence (31, 32) and increased resistance to SDS and hydrophobic antibiotics (22). We demonstrated that *tusA* deletion resulted in resistance to cationic drugs through the increased expression of *ompF* and *ompX* (Fig. 5). The outer membrane porins facilitate the passive transport of molecules, and the active export of drugs that have once entered the cell is unlikely. Additionally, cationic drugs exert antibacterial activity primarily by disrupting negatively charged cell membranes, without penetrating the cells (33). Therefore, we believe that the cationic drug resistance of Δ*tusA* is not because of a change in cationic drug permeability. This is consistent with the observation that the sensitivity of Δ*ompF* or Δ*ompX* to cationic drugs was comparable to that of the wild-type (Fig. 5D and E). Omp proteins and lipopolysaccharides form patches on the outer membrane, and their distribution and network determine the outer membrane properties (34). Increased expression of *ompF* and *ompX* may alter outer membrane properties, thereby conferring resistance to cationic drugs in Δ*tusA*.

## METHODS

### Bacteria and culture conditions

*E. coli* BW25113 and single deletion mutants were provided from the Japan National BioResource Project (BW25113, ME9062; Δ*tusA*, JW3435-KC; Δ*fur*, JW0669-KC; Δ*flhC*, JW1880-KC; Δ*rpoS*, JW5437-KC; Δ*ompX*, JW0799-KC; Δ*ompF*, JW0912-KC; and Δ*ompW*, JW1248-KC). The bacterial strains used in this study were cultured overnight on LB agar medium at 37 °C. The colonies were aerobically cultured in 5 mL LB liquid medium (1 % tryptone, 0.5 % yeast extract, and 1 % NaCl) or low-salt liquid medium (1 % tryptone and 0.25 % NaCl) in 50 mL Nunc conical sterile polypropylene centrifuge tubes (Thermo Fisher Scientific, US, MA) at 37 °C for 18 h. Double gene-knockout mutants were constructed by phage P1*vir*-mediated transduction (35) from the Δ*tusA* mutant into Keio knockout mutants in which the kanamycin resistance gene was deleted (36) as the recipient strains.

### RNA sequencing

Total RNA was extracted and sequenced as previously described (37). A 50 µL aliquot of overnight culture of BW25113 and Δ*tusA* was inoculated into 5 mL LB medium and cultured aerobically at 37 °C. When the OD600 of the culture reached 0.7, 1.8 mL of the culture was mixed with 200 µL of phenol in 5 % ethanol by vortexing, cooled in ice water for 5 min, and centrifuged (21,500 × g, 2 min). The cells were frozen in liquid nitrogen and stored at −80 °C for 2 h. The precipitate was dissolved in 200 µL of lysis buffer (TE buffer, 1 % lysozyme, and 1 % SDS) and incubated at 65 °C for 2 min. RNA was extracted from the samples using the RNeasy Minikit (Qiagen) following the manufacturer’s protocols. rRNA was removed using the NEBNext rRNA Depletion Kit, and the total RNA was converted to DNA libraries using the TruSeq Stranded Total RNA Kit (Illumina). RNA sequencing was performed using the NovaSeq 6000 system (Illumina) to generate 100-base pair-end reads of at least 4 GB per sample. RNA reads were mapped to the *E. coli* W3110 reference genome (NC_007779.1) using the CLC Genomics Workbench software (version 11.0). DEGseq2 analysis (38) was performed using the read count data for each gene and R software (version 4.4.2). Gene symbols were assigned to the locus tags using data from the Profiling of *E. coli* Chromosome database when they were not assigned during the CLC Genomics Workbench. The adjusted p-values and fold-change values of each gene were plotted using GraphPad Prism 9. Gene ontology enrichment analysis was performed using ShinyGO ver 0.77 with *E. coli* MG1655 (Taxonomy ID: 511145) as the reference data (39). Data for regulon analysis of differentially expressed genes were downloaded from regulonDB v13.5.0 (40).

### Immunofluorescence

To visualize the flagella, 100 µL of the bacterial culture was placed on a poly-L-lysine-coated cover glass and incubated for 1 h. After washing thrice with phosphate-buffered saline (PBS), 300 µL of anti-flagellin antibody (ab93713; Abcam, UK) diluted 1/4000 in PBS was added and incubated at 37 °C for 1 h. After washing with PBS thrice, 300 µL of Alexa Fluor 488 Donkey anti-rabbit IgG (406416; BioLegend, CA, U.S.A.) diluted 1/4000 with PBS was added to the plate and incubated at 37 °C for 40 min. After washing thrice with PBS, the flagella were observed using an Axiovert 5 fluorescence microscope equipped with an objective Plan-Apochromat 63x/1.4 Oil and Axiocam 305 color camera (Zeiss, Germany). The frequency of flagella was calculated by particle measurement using Fuji software v2.14 (41) for triplicate images captured randomly with identical excitation intensity and magnification.

### Swimming assay

To assess swimming motility, 1 µL of a bacterial culture in LB (OD_600_ = 5) was inoculated into LB or low salt medium containing 0.25 % agar. The swimming halo was photographed after 27 or 17 h of incubation at 37 °C, and its diameter was measured using Fiji software v2.14 (41).

### Assessment of cationic drug resistance

CTAB, CPC, and PS were dissolved in ultrapure water at concentrations of 50, 32.5, and 20 mg/mL, respectively, and added to autoclaved LB agar. SDS and Triton X-100 were dissolved in ultrapure water at 8 % and subsequently mixed with an equal amount of 2x concentrated autoclaved LB agar medium. *E. coli* cultures in LB were serially diluted 10-fold in LB broth in a 96-well microplate. Diluted samples (5 µL for LB control, CTAB, CPC, and PS or 3 µL for SDS and Triton X-100) were spotted onto LB plates using an 8-channel micropipette. The plates were incubated at 37 °C for 18 h and photographed using a digital camera.

## Acknowledgments

This work was supported by MEXT KAKENHI grants (Grant Nos. 22K19435, 23K24131, 22K14892, 24K21872, and 24K01760). We thank the Japan National Bioresource Project – *E. coli* (National Institute of Genetics, Japan) for providing the Keio collection.

---

**Table S1. RNAseq analysis data of wild-type BW25113 and Δ*tusA*.**

